# Differential mutational profile of SARS-CoV-2 proteins across deceased and asymptomatic patients

**DOI:** 10.1101/2021.03.31.437815

**Authors:** Rezwanuzzaman Laskar, Safdar Ali

## Abstract

The SARS-CoV-2 infection spread at an alarming rate with many places showed multiple peaks in incidence. Present study involves a total of 332 SARS-CoV-2 sequences from 114 Asymptomatic and 218 Deceased patients from twenty-one different countries. The mining of mutations was done using the GISAID CoVSurver (www.gisaid.org/epiflu-applications/covsurver-mutations-app) with the reference sequence ‘hCoV-19/Wuhan/WIV04/2019’ present in NCBI with Accession number NC-045512.2. The impact of the mutations on SARS-CoV-2 proteins mutation was predicted using PredictSNP1(loschmidt.chemi.muni.cz/predictsnp1) which is a meta-server integrating six predictor tools: SIFT, PhD-SNP, PolyPhen-1, PolyPhen-2, MAPP and SNAP. The iStable integrated server (predictor.nchu.edu.tw/iStable) was used to predict shifts in the protein stability due to mutations. A total of 372 variants were observed in the 332 SARS-CoV-2 sequences with several variants incident in multiple patients accounting for a total of 1596 incidences. Asymptomatic and Deceased specific mutants constituted 32% and 62% of the repertoire respectively indicating their exclusivity. However, the most prevalent mutations were those present in both. Though some parts of the genome are more variable than others but there was clear difference between incidence and prevalence. NSP3 with 68 variants had total occurrence of only 105 whereas Spike protein had 346 occurrences with just 66 variants. For Deleterious variants, NSP3 had the highest incidence of 25 followed by NSP2 (16), ORF3a (14) and N (14). Spike protein had just 7 Deleterious variants out of 66. Deceased patients have more Deleterious than Neutral variants as compared to the symptomatic ones. Further, it appears that the Deleterious variants which decrease protein stability are more significant in pathogenicity of SARS-CoV-2.

## Introduction

The causative agent of ongoing COVID-19 global pandemic is Severe Acute Respiratory Syndrome Coronavirus 2 (SARS-CoV-2) which belongs to family *Coronaviridae* characterized by single strand positive sense RNA genome (Sun et al., 2020). As of 31^st^ March 2021, there were 5,52,566, active cases, 1,14,34,301 discharged cases and 1,62,468 deaths in India due to SARS-CoV-2 https://www.mohfw.gov.in/). The same time, as per WHO there have been 27,349,248 confirmed cases of COVID-19, including 2,787,593 deaths worldwide due to COVID-19 (covid19.who.int).

As the SARS-CoV-2 infection spread at an alarming rate and many places showed multiple peaks in incidence, the virus has been replicating and accruing mutations in the process. There have been multiple reports to assess and understand the impact of these mutations (Chen et al., 2020; Li et al., 2020; van Dorp et al., 2020; Wang et al., 2020). These reports have focused on novel and recurrent variations and their impact on infectivity and antigenicity. However, the assessment of impact of mutations on SARS-CoV-2 is an ongoing process.

In our earlier study, we performed the mutational analysis of 611 genomes from India extracted n June 2020 and analyzed its impact on proteins. Therein we observed a differential mutation profile in viral genomes across deceased and asymptomatic patients (Laskar and Ali, 2021). However, one lacuna in the study was very limited number of samples as we had focused on India only but the indications therein formed the basis of present study from samples all over the world to ascertain the differential mutational profile of deceased and asymptomatic samples in order to understand the possible clinical implications of protein variants.

## Materials and Methods

### Sequence Congregation

On 10^th^ November 2020, we retrieved 6853 sequence from GISAID (www.gisaid.org) using the data filter ∼ virus name: hCoV-19 - Host: Human - Complete – High Coverage. Subsequently, they were screened for availability of clinical status which revealed 122 and 542 sequences from Asymptomatic and Deceased patients respectively. The final screening parameter applied was that of age wherein all samples of over 60 years were excluded. This was done because in deaths of people over 60 years, co-morbidities and other physiological aspects are involved (Omori et al., 2020).

Since our aim is to ascertain the impact of mutations, if any, on the deaths due to SARS-CoV-2, thus we excluded the samples where there are high chances of other factors contributing to death. Thereon, ‘a total of 332 SARS-CoV-2 sequences from 114 Asymptomatic and 218 Deceased patients were included in the study from twenty-one different countries. Details of the sequences used have been provided in Table 1 and Supplementary file 1. There were ten countries which had no asymptomatic patients and five countries with no deceased patients. These included 200 males, 72 females and 60 patients for which gender information was not available.

**Table 1:** Country-wise distribution of samples used in the study.

| **S No** | **Country** | **Total** | **Asymptomatic** | **Deceased** |
| --- | --- | --- | --- | --- |
| 01 | Bangladesh | 5 | 5 | 0 |
| 02 | Belgium | 2 | 1 | 1 |
| 03 | Brazil | 62 | 2 | 60 |
| 04 | Colombia | 3 | 0 | 3 |
| 05 | Costa Rica | 2 | 0 | 2 |
| 06 | Czech Republic | 3 | 1 | 2 |
| 07 | Dominican Republic | 3 | 3 | 0 |
| 08 | India | 68 | 28 | 40 |
| 09 | Indonesia | 2 | 0 | 2 |
| 10 | Italy | 3 | 1 | 2 |
| 11 | Japan | 59 | 59 | 0 |
| 12 | Kuwait | 2 | 2 | 0 |
| 13 | Lebanon | 1 | 0 | 1 |
| 14 | Mexico | 4 | 0 | 4 |
| 14 | Oman | 1 | 0 | 1 |
| 16 | Russia | 3 | 0 | 3 |
| 17 | Saudi Arabia | 70 | 0 | 70 |
| 18 | South Africa | 1 | 0 | 1 |
| 19 | Sri Lanka | 1 | 0 | 1 |
| 20 | Turkey | 8 | 8 | 0 |
| 21 | United States | 29 | 4 | 25 |
|  |  | **332** | **114** | **218** |

### Mining of Mutations in Proteins

The mining of non-synonymous mutations from the selected 332 sequences was done utilizing the GISAID CoVSurver (www.gisaid.org/epiflu-applications/covsurver-mutations-app) web resources for sequence variance analysis. Sequence datasets were aligned with the reference sequence ‘ hCoV-19/Wuhan/WIV04/2019’. The reference amino acid sequence of each protein for further analysis was downloaded from the same server. The reference sequence is hundred percent identical to the NCBI sequence with Accession number NC-045512.2 from Wuhan, China. It has been used as reference for our earlier study on SARS-CoV-2 genomes from India (Laskar and Ali, 2021, 2020).

### Pathogenicity Prediction of Mutations

The impact of the mutations on SARS-CoV-2 proteins mutation was estimated using PredictSNP1(loschmidt.chemi.muni.cz/predictsnp1) which is a meta-server integrating six predictor tools: SIFT, PhD-SNP, PolyPhen-1, PolyPhen-2, MAPP and SNAP. The SNAP and PhD-SNP tools are based on supervised machine learning algorithms that have been trained in massive datasets to “learn” to distinguish between pathogenic and benign variants. The protein sequence and structure method predict how mutations influence the protein phenotype on the basis of the SNP position in the protein structure behind the PolyPhen-1 and PolyPhen-2 predictors. The MAPP and SIFT tools, on the other hand, determine the pathogenicity of mutations based on the conservation of specific amino acids by sequence and evolutionary conservation methods across various species. PredictSNP extracted the individual score of these tools to homogenize the assessment and generate its own confidence score as a percentage of expected accuracy, ranging from 0 to 100 per cent. It has also designed three distinct datasets like PMD, MMP and OVERFIT in order to eliminate bias, duplicity and inconsistency (Bendl et al., 2014). Subsequently, the variants are categorized as ‘ Neutral’or ‘ Deleterious’, and these predictions are more robust and accurate than the prediction provided by any individual predictor.

### Stability Shifts of Protein Mutations

The iStable integrated server (predictor.nchu.edu.tw/iStable) encompasses two machines learning based predictor iMutant and MUpro along with thermodynamics parameters in order to predict shifts in the protein stability related to mutations. i-Mutant, an SVM-based tool, predicts protein stability attributable to mutant form in free energy change value [DDG=DG(NewProtein)-DG(WildType) in Kcal/mol]. It classifies the variant in terms of decreasing (DDG<0) or increasing (DDG>0) protein stability. MUpro, an SVM and neural network-based tool, evaluate the mutational impact of protein stability on predictive confidence score (Conf. Score). The score varies from -1 to 1, with Conf. Score <0 signifies a decrease in stability and Conf. Score >0 as an increase in protein stability. iStable utilizes sequence information and predicts the meta-result as an increase or decrease in stability in terms of confident score where a higher score indicates more confidence in the prediction (Chen et al., 2013).

## Results and Discussion

### Mutation Incidence and Prevalence

A total of 372 variants were observed in 332 SARS-CoV-2 sequences with several variants being incident in multiple patients accounting for a total of 1596 variants. The distribution of variants across different proteins of SARS-CoV-2 have been shown in Figure 1, Table 2 and Supplementary file 2. The distribution and incidence of variants can be analyzed through several aspects.

**Table 2:** Distribution of Deleterious (Red) and Neutral (Black) mutations along with D/N ratio (Deleterious/Neutral) across different proteins of SARS-CoV-2 in Deceased (De) and Asymptomatic (As) patients.

| S No | Protein | Total Variations (De+As+Bo) | Deceased (De) | Asymptomatic (As) | Both (Bo) | Deceased (De) | D/N Ratio | Asymptomatic (As) | D/N Ratio | Both (Bo) | D/N Ratio |
| --- | --- | --- | --- | --- | --- | --- | --- | --- | --- | --- | --- |
| 1 | E | 6(4+2+0) | C43F<br>L21F; T30I; P71S | R38C<br>I46V | Nil | 1, 3 | 0.33 | 1, 1 | 1 | 0, 0 | NA |
| 2 | M | 7(5+2+0) | A2V; D3G; L29F;<br>V66L; H125Y | A98S; C33F | Nil | 0, 5 | 0 | 0, 2 | 0 | 0, 0 | NA |
| 3 | N | 29(12+13+4) | A119S; S193I;<br>S194L; T247I;<br>K373N; D402Y;<br>Q418H<br>R41L; L139F;<br>H145Y; S327L;<br>K374N | D225Y; Q9H;<br>R195I; S197L;<br>S202N<br>Q7K; D103N;<br>I351V; P199S;<br>H300Y; T334A;<br>T379I; A397V | G204R;<br>P13L<br>R203K;<br>I292T | 7, 5 | 1.4 | 5, 8 | 0.625 | 2, 2 | 1 |
| 4 | ORF3a | 20(13+5+2) | S40L; L41H;<br>A51S; I62M;<br>A110V; R126S;<br>G172V; Q185K;<br>G251V; S253P<br>I118T; M125I;<br>V225F | T89I; G196V;<br>G224C<br>L15F; L52I | Q57H<br>S171L | 10, 3 | 3.33 | 3, 2 | 1.5 | 1, 1 | 1 |
| 5 | ORF6 | 5(3+1+1) | I11R; R20S; W27L | V5F | I33T | 3, 0 | ∞ | 1, 0 | ∞ | 1, 0 | ∞ |
| 6 | ORF7a | 8(6+2+0) | V74F; S83L;<br>V93F; Q94E<br>A8S; V108L | T39I<br>A105V | Nil | 4, 2 | 2 | 1, 1 | 1 | 0, 0 | NA |
| 7 | ORF8 | 9(5+2+2) | Q23H<br>D34G; A51S;<br>A51V; A65D | D119G; V99L | L84S;<br>S24L | 1, 4 | 0.25 | 0, 2 | 0 | 0, 2 | 0 |
| 8 | NSP1 | 5(1+4+0) | R124C | R24C; D75N;<br>I114T<br>E57K | Nil | 0, 1 | 0 | 3, 1 | 3 | 0, 0 | NA |
| 9 | NSP2 | 27(13+12+2) | D23Y; E66G;<br>P181T; S211F;<br>T371I; Q496P<br>I21V; P129L;<br>S196L; A247V;<br>R380C; T429I;<br>T434I | R27C; K110N;<br>M141T; A159V;<br>G339S; K347N;<br>Q427H; L520F;<br>V577F<br>I120F; S430L;<br>V198I; | T85I<br>A318V | 6, 7 | 0.86 | 9, 3 | 3 | 1, 1 | 1 |
| 10 | NSP3 | 68(40+26+2) | D112Y; D178Y;<br>A256V; G337C;<br>L425P; T428I;<br>P466L; L623I;<br>L689F; R810C;<br>L862F; K1083I;<br>D1121G; L1244F;<br>G1476S; A1914D<br>A1S; S126L;<br>P200S; G212S;<br>P389L; V481L;<br>K487N; A496V;<br>T504P; S716I;<br>T724I; P822L;<br>K1077N; K1211N;<br>P1228L; S1314G;<br>A1324V; S1424P;<br>A1431V; S1670F;<br>S1682F; K1693N;<br>V1795I; A1872V | S1197R; D163Y;<br>F430S; G470S;<br>G470V; T779I;<br>K945N; V1768G;<br>S1807F<br>P153S; I341M;<br>K379E; K412N;<br>I458T; A480V;<br>A488T; M494I;<br>P723S; A994D;<br>K1042N; T1072I;<br>C1114I; P1132S;<br>M1278K; D1674N;<br>V1762A | A861V;<br>T1198K | 16, 24 | 0.67 | 9, 17 | 0.53 | 0, 2 | 0 |
| 11 | NSP4 | 14(10+3+1) | S148P; G309C<br>W6C; T204I;<br>T214I; A231V;<br>T295I; M366I;<br>A457V; S59P | G196D; F308Y<br>M33I | A307V | 2, 8 | 0.25 | 2, 1 | 2 | 0, 1 | 0 |
| 12 | NSP5 | 11(11+0+0) | A266V; L89F;<br>T196M; V18I<br>K12R; G15S; T45I;<br>G71S; P96S;<br>V125I; T198I | Nil | Nil | 4, 7 | 0.57 | 0, 0 | NA | 0, 0 | NA |
| 13 | NSP6 | 9(7+2+0) | V190F; I162T<br>A88V; L125F;<br>V149F; Y153F;<br>M183I | C113F; L37F | Nil | 2, 5 | 0.4 | 0, 2 | 0 | 0, 0 | NA |
| 14 | NSP7 | 2(2+0+0) | L71F; S25L | Nil | Nil | 2, 0 | ∞ | 0, 0 | NA | 0, 0 | NA |
| 15 | NSP8 | 4(2+2+0) | A74V<br>T123I | G144S; V26I | Nil | 1, 1 | 1 | 0, 2 | 0 | 0, 0 | NA |
| 16 | NSP9 | 1(0+1+0) | Nil | L42F | Nil | 0, 0 | NA | 0, 1 | 0 | 0, 0 | NA |
| 17 | NSP10 | 3(2+1+0) | K124R; T111I | T101I | Nil | 0, 2 | 0 | 0, 1 | 0 | 0, 0 | NA |
| 18 | NSP12 | 21(17+1+3) | D140G; T248I;<br>L308M; V354A;<br>V354G; S814P<br>L8F; N9K; M196I;<br>P227L; A250V;<br>T252N; L329I;<br>N491S; V880I;<br>A878S; H928Q | V354L | A97V;<br>L638F<br>P323L | 6, 11 | 0.55 | 1, 0 | ∞ | 2, 1 | 2 |
| 19 | NSP13 | 18(7+10+1) | S74P; T115I<br>L297F; D466N;<br>T501I; I258V;<br>I334V | P47S; A237T;<br>S485L; P504L;<br>Y541C; M576I<br>G54S; S236I;<br>K524N; A553V | S166A | 2, 5 | 0.4 | 6, 4 | 1.5 | 0, 1 | ∞ |
| 20 | NSP14 | 17(14+3+0) | S28I; R163C;<br>L177F; T215I<br>N3H; K32R;<br>P140S; I45K;<br>V161I; C216F;<br>A320V; K349N;<br>Q354H; T372I | D345G; H26Y;<br>T206I | Nil | 4, 10 | 0.4 | 0, 3 | 0 | 0, 0 | NA |
| 21 | NSP15 | 14(10+4+0) | S287L; Y225F<br>T33I; V38F;<br>A171V; G129S;<br>E223A; K259N;<br>E260K; P270T; | I143N; R206S<br>V172L; S261L | Nil | 2, 8 | 0.25 | 2, 2 | 1 | 0, 0 | NA |
| 22 | NSP16 | 8(4+3+1) | T117I; I267T<br>A116V; K160R | K263N<br>M17I; T35I | S166A | 2, 2 | 1 | 1, 2 | 0.5 | 0, 1 | 0 |
| 23 | Spike | 66(43+20+3) | C136G; Y489H;<br>S939F<br>P9L; S12F; T22I;<br>A27S; L54F; F59S;<br>A67V; R78M;<br>Y144F; M153I;<br>H146Q; Q183H;<br>V213A; A264S;<br>T299I; V382L;<br>T478K; E484K;<br>S494P; N501Y;<br>T572I; D574Y;<br>Q613H; A647S;<br>A688S; A688V;<br>T791I; P812L;<br>A879S; N978D;<br>L1063F; T1117I;<br>G1124V; D1153Y;<br>P1162S; V1176F;<br>N1192K; I1225T;<br>M1237L; I870V | S46L; D88H;<br>G769V; D1084Y<br>D627P; G181V;<br>G476S; I119V;<br>K1086R; K77M;<br>K854N; M1229I;<br>N679K; P1079T;<br>P681R; Q218H;<br>R765L; T29I;<br>V1228L; V47I | D614G;<br>E583D;<br>L5F | 3, 40 | 0.075 | 4, 16 | 0.25 | 0, 3 | 0 |

**Figure 1:**
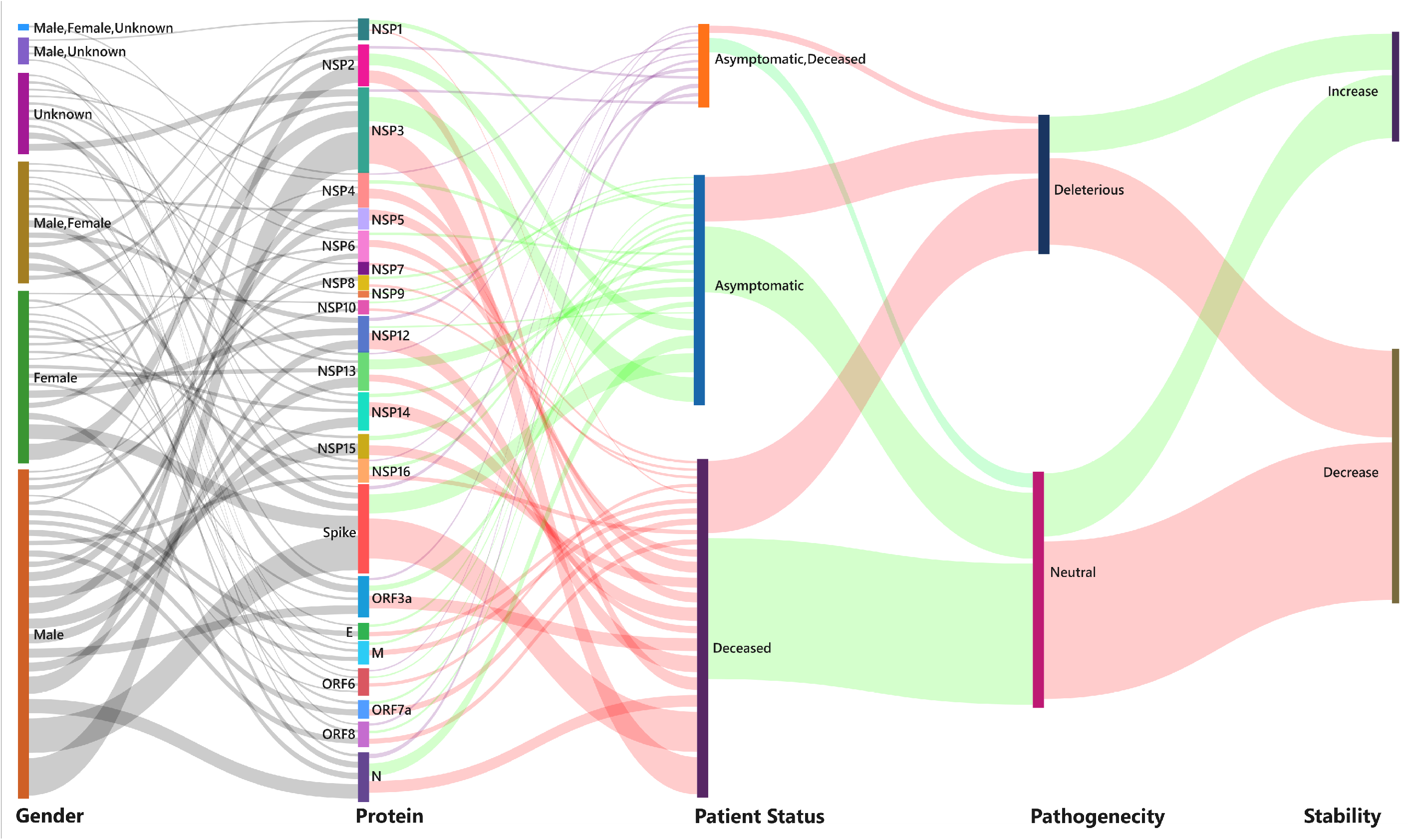
Distribution of variant sites of SARS-CoV-2 proteins across Gender; Patient status (Deceased/Asymptomatic); Pathogenicity (Deleterious/Neutral) and Stability.

First, the incidence of mutations with respect to gender as summarized in Fig 2A. The patients for whom gender information wasn’t available have been mentioned as unknown. There were 45 mutations (12%) present in both males and females but these variants were the most prevalent ones as they accounted for around 69% of the total incidences. Also, the male and female specific 221 and 75 mutations (59% and 20%) accounted for a meagre 17% and 5% respectively of total incidence. Partly this difference can be attributed to the skewed sample set in favor of the males. Klein and Flanagan have reviewed the studies assessing the variable nature of immune responses in males and females (Klein and Flanagan, 2016) but since the variants most incident are present in both genders in our study and assuming the variations can potentially alter the immune response, the chances of it in a gender-dependent manner seems rare.

**Figure 2:**
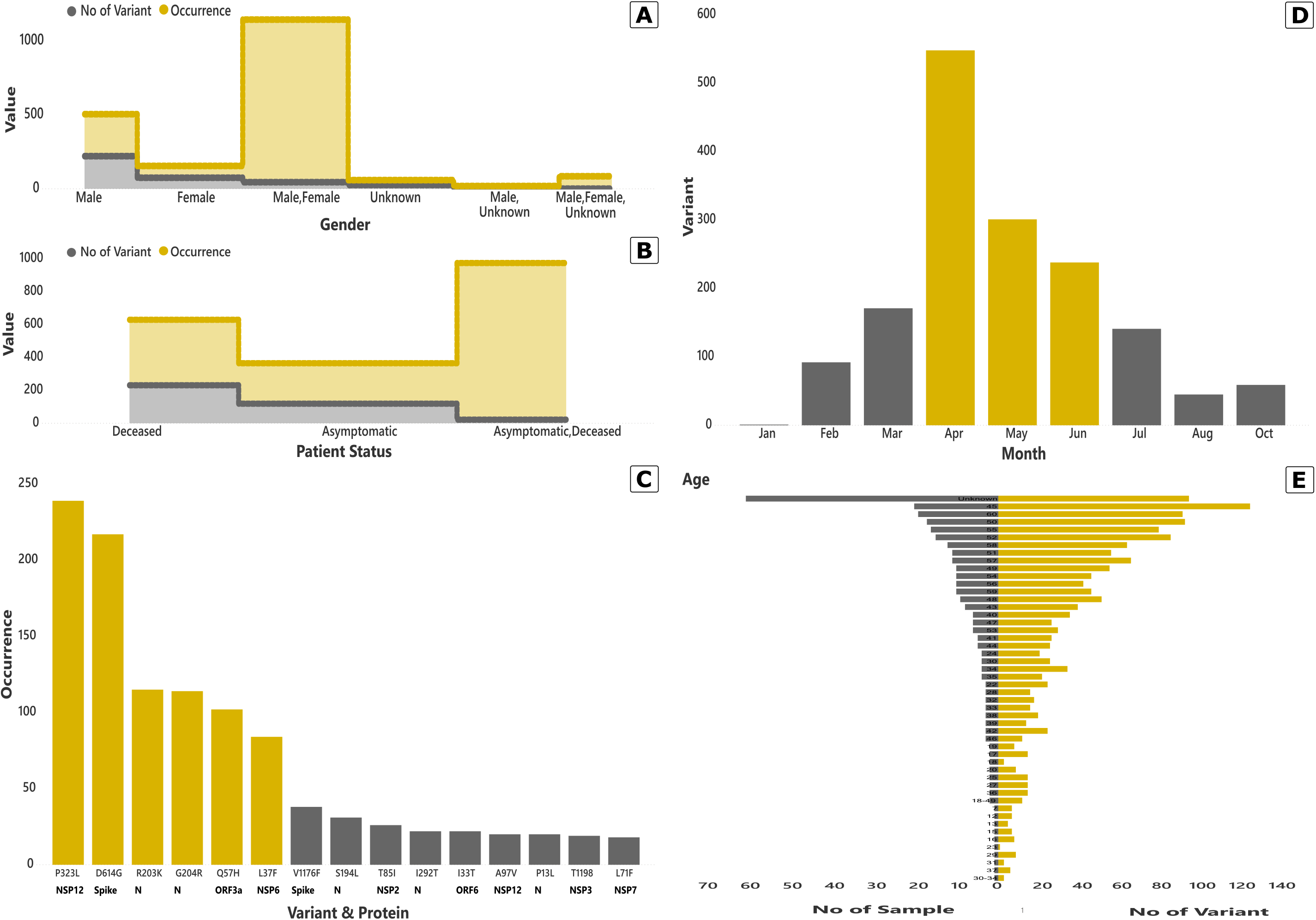
Details of mutations of SARS-CoV-2 proteins. **A)** Gender wise distribution of variants and its prevalence in studied genomes. **B)** Distribution of observed variants according to clinical status of patients as per Deceased/Asymptomatic/Both **C)** Most prevalent variants across proteins observed in the study. **D)** Timeline of incidence of observed variations. **E)** Age wise distribution of samples of the present study.

Though there 60 patients with unknown gender, the fact that overall distribution of mutants and incidence in the studied population is around 90% for males, females or both means the data from unknown will not have much of an impact on gender wise distribution and incidence of mutants. Almost all of these 60 patients are asymptomatic and belong to the Diamond Princess cruise ship from Japan which has become benchmark study about how wearing masks reduces viral load and hence even if the person gets infected chances of it being mild or asymptomatic are enhanced substantially (Batista et al., 2020; Gandhi et al., 2020).

Subsequently, we moved to the second aspect of analysis: patient status. The question we wanted to address was are any mutations specific to the deceased or asymptomatic patients? The distribution and incidence of mutations as per patient status has been shown in Figure 2B and Supplementary file 2. Interestingly, 22 mutations present in both deceased and asymptomatic accounted for 952 (60%) of the total occurrence of mutations. Asymptomatic specific mutants encompassing 119 variants (32%) accounted for only 246 (15%) of the total observed mutations whereas the corresponding values for Deceased specific mutants was 231 (62%) variants with 398 (24%) occurrences respectively. The details of these mutations have been provided in Table 2 and Supplementary file 2 and their impact on proteins are discussed later.

Evidently, some mutations are getting represented at a much higher rate as compared to others. The fifteen most prevalent mutations along with their incident frequency and localization have been shown in Figure 2C. P323L (NSP12) is the most prevalent variant present followed by D614G (Spike protein) with over two hundred incidences each. Most of these prevalent variants are common to both males and females, which is expected as per gender wise distribution data.

The timeline for accumulation of mutants has been shown in Figure 2D and evidently the maximum number of mutations accrued in April 2020 wherein the incidence of COVID-19 was at its peak. The age wise distribution of mutants has been shown in Figure 2E. An absolute correlation between age and incidence can’t be drawn due to uneven representation of all age groups. Furthermore, since the samples are from different countries and the virus is behaving differently across geographical locations, the data has to be interpreted with caution. Thereby, looking at the mutation incidence with reference to countries seemed rational.

We analyzed the data in terms of number of mutants per sample for each country as represented in Figure 3 and Supplementary file 4. Saudi Arabia with maximal representation of 70 samples in the study was contributing 3.93 variants per sample which is amongst the lowest in the group. Contrastingly, Bangladesh with highest value of 10 variants per sample is being represented by just 5 samples in the study. Thus, its explicit that some nations are exhibiting more variations in SARS-CoV-2 as compared to others. Differential health and gene profile of individuals might be factors contributing to it. We also looked at the variant per sample data for deceased and asymptomatic samples respectively for each country as shown in Supplementary file 4. Bangladesh had the highest variants per sample of 10 followed by India (6.36) for asymptomatic patients whereas it was highest of 9.5 for Indonesia followed by Brazil (6.75) for deceased patients. The fact that some countries had no representation in asymptomatic or deceased samples makes any direct inference plausible but a geographical evolution of the virus is surely supported by the observed data.

**Figure 3:**
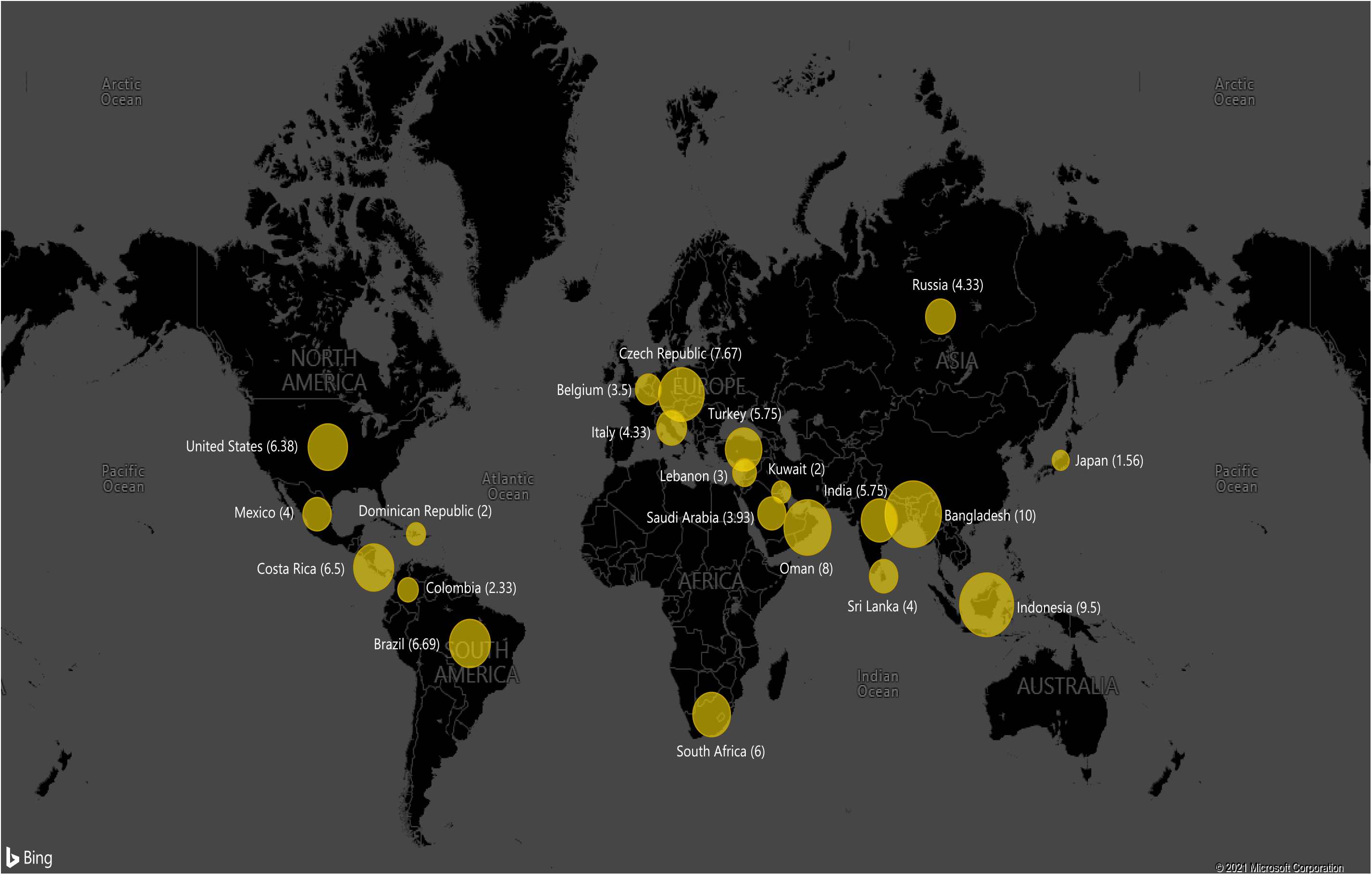
Country wise distribution of variants of SARS-CoV-2 proteins.

### Impact of Mutations on Proteins of SARS-CoV-2

The impact of mutations on SARS-CoV-2 proteins was assessed using three aspects as highlighted in Figure 4. These include distribution of variants across SARS-CoV-2 proteins; pathogenicity of the variants in term of deleterious or neutral and assessing the stability of proteins in presence of variants.

**Figure 4:**
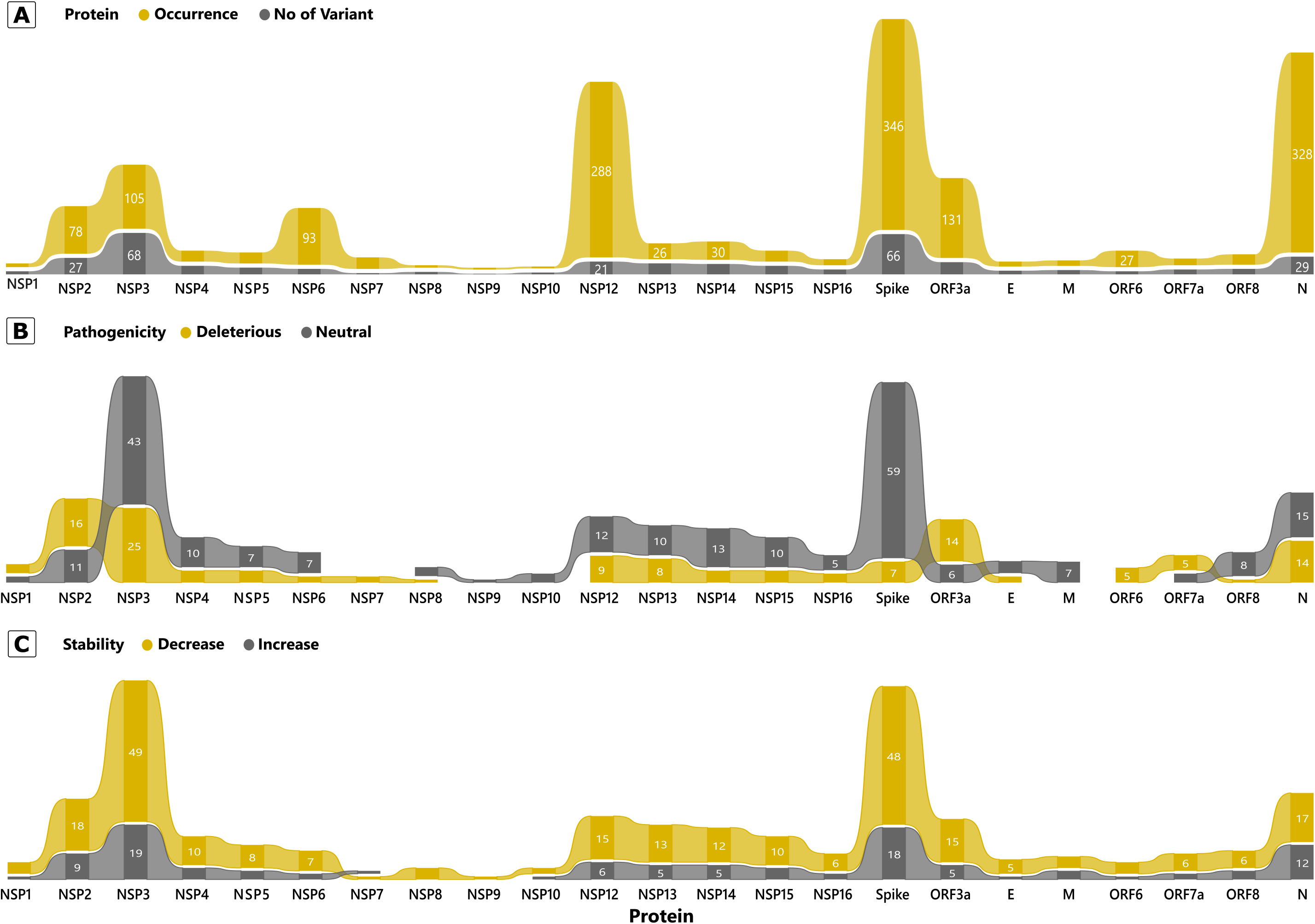
Mutation incidence and its impact on SARS-CoV-2 proteins. **A)** Number of variants and its prevalence in studied genomes. **B)** Pathogenicity Prediction of the variants in terms of Deleterious or Neutral. **C)** Stability shift prediction of the observed variants.

The distribution of variants across different SARS-CoV-2 proteins revealed some interesting observations. First, variants incidence in some proteins was relatively higher than others as evident in Figure 4a and Supplementary file 2. NSP3 housed a maximum of 68 variants closely followed by Spike protein with 66 variants. N protein and NSP2 with 29 and 27 variants respectively were a distant third and fourth. Thus, some parts of the genome are more variable whereas others are more conserved. These have led to emergence of novel variants around the world which have been so far without clinical manifestations but the situation might change any moment. Secondly, there was clear difference between incidence and prevalence. NSP3 with 68 variants had total occurrence of only 105 whereas Spike protein had 346 occurrences (66 variants). Comparatively, N protein and NSP12 had 328 (29 variants) and 288 (21 variants) occurrences in the study. This clearly implies that clustering of mutations doesn’t imply higher prevalence.

In order to ascertain the significance of differential incidence and prevalence, we analyzed the pathogenicity of the variants as Deleterious or Neutral which has been shown in Figure 4b, Table 2 and Supplementary file 3. Their presence across Asymptomatic and Deceased samples across different countries has been discussed in the next section. The basic premise for the study was that a protein having more variants will be contributing to the viral evolution only if they are Deleterious. Neutral mutations wouldn’t be affecting the protein per se. In terms of incidence of Deleterious variants, NSP3 had the highest of 25 followed by NSP2 (16), ORF3a (14) and N (14). Spike protein had just 7 Deleterious variants out of 66 which partly explains that despite of so many variants it hasn’t much impacted the viral pathogenesis yet. It is also interesting to note that only four proteins NSP1, NSP2, ORF3a and ORF7a had more Deleterious than Neutral mutants. Further, NSP7 and ORF6 had only Deleterious mutants (Figure 4b, Table 2). Thus, we can say that even a protein with very few mutations can be a more determining factor in viral evolution.

Assessment of the protein stability in lieu of mutations was thereby ascertained and has been represented in Figure 4c. Majority of variants resulted in a decrease in protein stability across all proteins. The three proteins which had maximum variants which increased protein stability included NSP3 (19), Spike (18) and N (12). Also, NSP3 (49) and Spike (48) had highest number of variants which decreased protein stability. The impact of these variants discussed individually is significant but their occurrence in population isn’t in isolation and hence the cumulative impact of mutations incident together is also required.

### Differential Mutational Profile Across Asymptomatic and Deceased Samples

The distribution of Deleterious and Neutral mutations across asymptomatic and deceased patients has been shown in Table 2 and Supplementary file 3. There are 12 proteins in which no variant was present in both asymptomatic and deceased samples suggesting a correlation between mutational status and disease profile. N protein had the highest of 4 variants across both sample sets. Moreover, except for N protein, NSP1, NSP3, NSP9 and NSP14, all other proteins had more variants associated with deceased patients as compared to asymptomatic ones. NSP5 and NSP7 had no mutations from asymptomatic patients whereas only NSP9 had nor variants coming from deceased patients.

Subsequently we analyzed the ratio of Deleterious is to Neutral mutations across asymptomatic and deceased samples as shown in Table 2. Most proteins had a higher D/N ratio for deceased samples indicating the implication of Deleterious mutations therein. The highest D/N ratio was 3.33 for OF3a followed by 1.4 for N protein. The fact that deceased patients have more Deleterious than Neutral variants as compared to the symptomatic ones is suggestive of these mutations being correlated to the disease status of the samples.

To further construe the link between the deceased host and the non-synonymous mutation, we analyzed the mutated protein clusters as follows: From Deceased Individuals (De) with Deleterious (D) pathogenicity and Increase in Stability (IS) and From Deceased Individuals (De) with Neutral (N) pathogenicity and Decrease in Stability (DS). There details have been shown in Table 3 and Supplementary file 3. When we analyzed mutations only from deceased patients and with deleterious pathogenicity, we observed that there were nine proteins in which all such mutations had decreasing stability suggesting the possibility of the decreased stability contributing to enhanced pathogenicity. Moreover, there were ten proteins which had mutations with increasing as well as decreasing stability. Amongst these there was on N protein which had four variants with increasing stability as compared to three with decreasing stability prediction. All other proteins had more stability decreasing variants as compared to increasing ones. Thus, we can say that the variants with decreasing stability are more significant in pathogenicity of SARS-CoV-2.

**Table 3:**
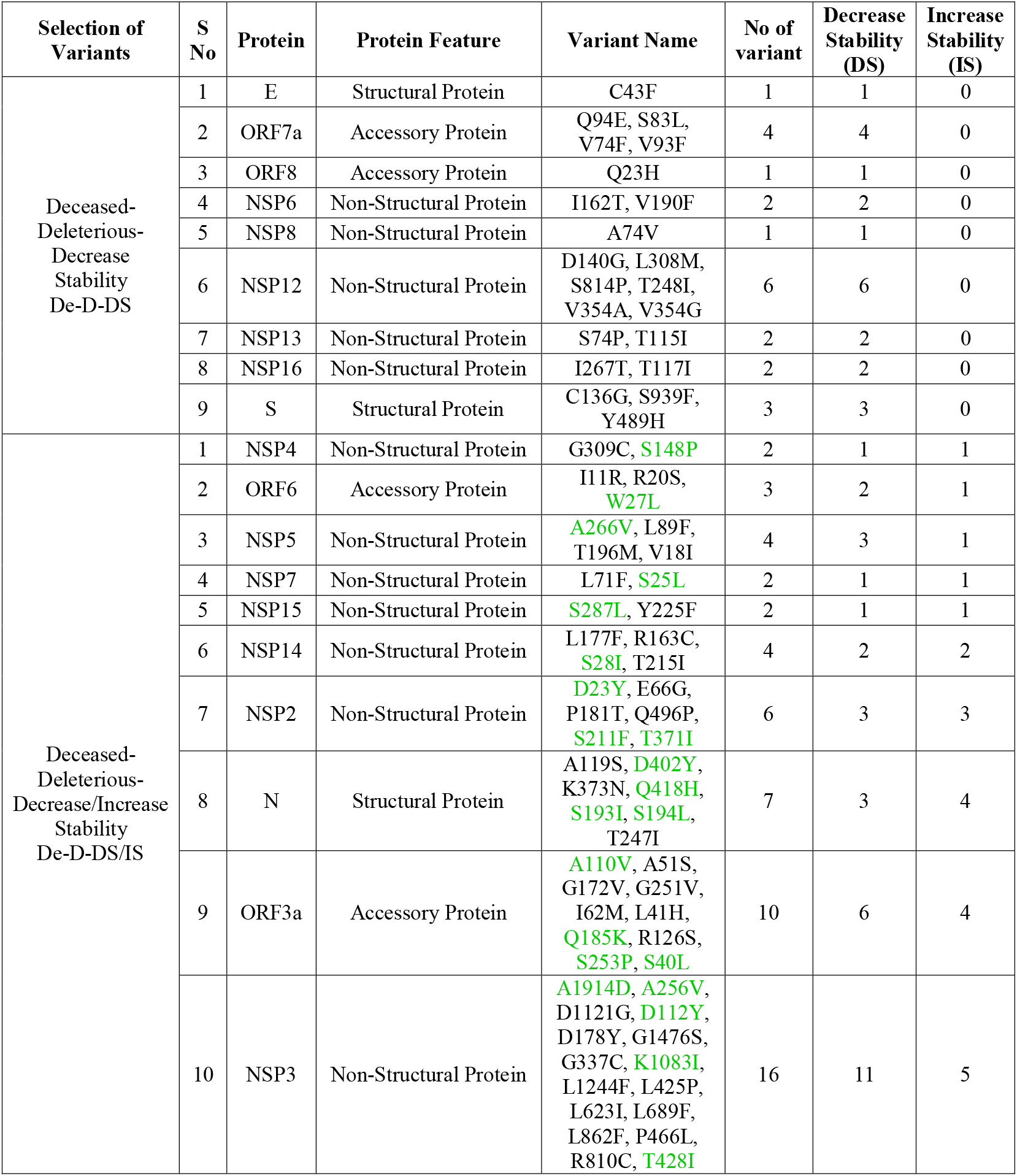
Correlation between Deleterious mutations from Deceased individuals and Protein Stability. *(Variants with increased stability have been shown in green)*

## Conclusions

A total of 372 variants were observed in 332 SARS-CoV-2 sequences with several variants being incident in multiple patients accounting for a total of 1596 variants. The 22 mutations present in both deceased and asymptomatic accounted for 60% of the total occurrence of mutations. Asymptomatic specific 119 variants accounted for 15% whereas Deceased specific 231 variants represented 24% of total occurrences respectively. Since, some countries had no representation in asymptomatic or deceased samples, inference about geographical correlation isn’t plausible from present dataset. Four proteins NSP1, NSP2, ORF3a and ORF7a had more Deleterious than Neutral mutants. When looking at deceased patients with deleterious pathogenicity prediction mutants, there were nine proteins in which all such mutations had decreasing stability suggesting the possibility of the decreased stability contributing to enhanced pathogenicity.

## Acknowledgements

The authors thank the Department of Biological Sciences, Aliah University, Kolkata, India for all the financial and infrastructural support provided. Authors acknowledge all the authors associated with originating and submitting laboratories of the sequences from GISAID’s EpiFlu™ (www.gisaid.org) Database on which this research is based.

## Declarations

### Competing Interests

The authors declare they have no competing interests.

### Ethics approval

Not Applicable.

### Availability of Data and Materials

All data pertaining to the study has been provided as Supplementary material of the manuscript.

### Authors’contributions

RL: Methodology, Investigation, Formal Analysis and Validation

SA: Conceptualization, Supervision, Formal Analysis and Writing.

## Supplementary files

Supplementary File 1: Details of asymptomatic samples used in the study (GISAID Reference; Age; Gender; Country)

Supplementary file 2: Localization of all the mutations observed in studied SARS-CoV-2 genomes

Supplementary files 3: Localization of variants affecting amino acid sequence in studied SARS-CoV-2 genomes (Distribution; Deceased/Asymptomatic; Deleterious/Neutral; Incidence; Stability)

Supplementary file 4: Distribution of samples and variants in the study

## Notes

### Competing Interest Statement

The authors have declared no competing interest.

